# Optimised multiplex amplicon sequencing for mutation identification using the MinION nanopore sequencer

**DOI:** 10.1101/2021.09.21.461312

**Authors:** Whitney Whitford, Victoria Hawkins, Kriebashne Moodley, Matthew J. Grant, Klaus Lehnert, Russell G. Snell, Jessie C. Jacobsen

## Abstract

**Objective:** Rapid, cost-effective identification of genetic variants in small candiate genomic regions remains a challenge, particularly for less well equipped or lower throughput laboratories. Application of Oxford Nanopore Technologies’ MinION sequencer has the potential to fulfil this requirement. We have developed a multiplexing assay which pools PCR amplicons for MinION sequencing to enable sequencing of multiple templates from multiple individuals which could be applied to gene-targeted diagnostics.

**Methods:** A combined strategy of barcoding and sample pooling was developed for simultaneous multiplex MinION sequencing of 100 PCR amplicons, spanning 30 loci in DNA isolated from 82 neurodevelopmental cases and family members. The target regions were chosen for further interegation because a potentially disease-causative variants had been identified in affected individuals by Illumina exome sequencing. The pooled MinION sequences were deconvoluted by aligning to custom references using the guppy aligner software.

**Results:** Our multiplexing approach produced interpretable and expected sequence from 29 of the 30 targeted genetic loci. The sequence variant which was not correctly resolved in the MinION sequence was adjacent to a five nucleotide homopolymer. It is already known that homopolymers present a resolution problem with the MinION approach. Interstingly despite equimolar quantities of PCR amplicon pooled for sequencing, significant variation in the depth of coverage (139x – 21,499x; mean = 9,050, std err = 538.21) was observed. We observed independent relationships between depth of coverage and target length, and depth of coverage and GC content. These relationships demonstrate biases of the MinION sequencer for longer templates and those with lower GC content.

**Conclusion:** We demonstrate an efficient approach for variant discovery or confirmation from short DNA templates using the MinION sequencing device. With less than 140x depth of coverage required for accurate genotyping, the methodology described here allows for rapid highly multiplexed targeted sequencing of large numbers of samples in a minimally equipped laboratory.

## Introduction

The rapid evolution of the study of genetic variation, and the discovery of disease susceptibility and causal variants has led to a demand for methods for rapid variant discovery or accurate verification. Identifying a causative disease mutation can guide clinical management plans and lead to the identification of life-saving treatments [1, 2]. The development of short read high throughput re-sequencing, particularly whole exome sequencing (WES) and whole genome sequencing (WGS) has accelerated this discovery process.

Decreasing cost has resulted in a rapid adoption of these re-sequencing techniques and an increase in the rate of new pathogenic mutations discovery [3, 4]. Trio sequencing (where the proband and their parents are sequenced) aids in variant analysis by determining the inheritance of putative causative variants. However, sequencing all members of a trio rather than just the proband is a significant extra cost. Sanger sequencing of selected PCR amplicons in close relatives such as parents is most often used to confirm inheritance of variants [2]. However, if there are numerous loci of interest in an individual or across a cohort, this can be a slow and costly exercise [5].

As an alternative to single amplicon, sample by sample, Sanger sequencing, we considered that there may be efficiency gains to be made by multiplex sequencing using the Oxford Nanopore Technologies (ONT) MinION nanopore sequencer. This device is a low-cost USB-powered sequencer that returns sequencing results in real-time [6]. Due to the low cost and small size of the MinION, it is a viable, easy to operate tool for genetics labs. The main limitation of MinION sequencing is the high nucleotide error rate [7], currently between ~5% and 15% [8]. However, the high error rate can be mitigated by using multiple reads to establish a consensus following alignment or assembly.

The barcoding of PCR amplicons with barcodes available from ONT enables simultaneous sequencing of multiple samples, reducing the per-sample cost. The effectiveness of sequencing multiplex PCR amplicons from multiple individuals has been demonstrated in forensics [9, 10], heritable genetic disease [11] and pharmacogenetic screening applications [12]. Notably, this approach played a significant role in tracking viral spread during the Zika [13] and COVID-19 [14-16] outbreaks. A previous report by Liou, et al. (2020) leveraged MinION sequencing as an alternative to Sanger sequencing for multilocus sequencing to strain type samples of *Staphylococcus aureus*, sequencing a pool of 672 PCR amplicons from the same sample [17]. This approach was assisted by a dual barcoding approach whereby barcodes (additional to the ONT native barcode) were incorporated into the PCR amplicons using primers with barcode sequence incorporated to the 5’ end of primers. This approach enabled the diferentiation of sequence of the same target region from multiple samples (in this case, typing 96 bacteria in the same sequencing run).

In this report, we present a method to pool different amplicons under ONT native barcodes to simultaneously sequence multiple amplicons, exceeding the number of barcodes used. We sequenced 100 total PCR amplicons from 82 participants pooled under ten barcodes, looking for sequence variance at 30 genetic loci. To our knowledge, this is the first report describing a method where diverse PCR amplicons can be pooled under the ONT native barcodes, allowing for high-thoughput sequencing of any amplicon using the MinION.

## Materials and methods

### Samples

The study cohort consisted of 82 people comprised of 55 individuals with autism spectrum disorder who had been whole exome sequenced (Supplementary Methods), and 27 parents of the cases who had not undergone high-throughput sequencing. The genetic variants selected for further investigation in the parents were considered to be putative disease-causing mutations (Supplementary Methods). These variants consisted of 28 single and two indels (a 1 bp deletion and a 2 bp insertion) (Supplementary Methods). DNA from the cohort was extracted from whole blood using the Qiagen Gentra Puregene Blood Kit (Hilden, Germany) or from saliva Oragene prepIT L2P saliva extraction kit (DNA GEnotek Inc, Ottawa, Canada) according to the manufacturer’s protocols.

### Individual PCR

Amplicons incorporating each of the putative mutations ranged in size from 122-707 bp (see Supplementary Table 1). Each PCR reaction mix consisted of 0.5 μM each of the forward and reverse primer, 1 × KAP2G Buffer A, 0.2 mM dNTP mix, and 0.5 units KAPA2G Robust DNA Polymerase, combined in a 25 μl reaction (Kapa Biosystems, Massachusetts, USA). The PCR cycling conditions were as follows: initial denaturation of 95°C for 3 min, followed by 35 cycles of 95°C for 15 s, 60°C for 15 s, and 72°C for 30 s, followed by a final 72°C extension for 1 min.

The PCR amplicons were purified using Ampure XP magnetic beads (Beckman Coulter, California, USA) and quantified by Qubit Fluorometric Quantification Broad Range Assay (Invitrogen, California, USA). All PCR amplicons were Sanger sequenced and the target variants identified for comparison.

### Sample Pooling and Barcoding

In order to analyse the MinION pooled sequence, a strategy for deconvoluting the sequence reads and assign a read uniquely to a single sample of origin is required. Thus, we leveraged the step of sequence alignment to separate the PCR amplicons pooled under a single barcode. Each demultiplexed barcoded pooled sequence was aligned to a barcode-specific customgenerated reference - specifically, a fasta file with the sequence of each PCR amplicon target region pooled within a specific barcode. As the deconvolution relies upon the assignment of sequence reads to the correct target sequence, similarity between target sequences within the same reference could result in reads aligning to the wrong target. In order to ensure the correct assignment of reads to the corresponding target reference sequence, PCR amplicons pooled under a single barcode should not contain significant sequence similarity.

The PCR amplicons in this experiment were divided into ten pools (ten DNA templates in each pool) with no pool containing more than one subject PCR amplicon to the same target region. As a result, there was no similarity within pooled PCR amplicons within individual barcodes. Similarity was determined by BLAST Global Alignment [17], where a fasta file of the target sequences for a single barcode was queried against itself using default parameters. For each barcode, each target region shared 0% similarity with all other targets.

0.16 pmol of each sample was added to each pool for a total of 1.6 pmol per pool. DNA templates within each pool were ligated to a specific barcode according to the 1D Native barcoding DNA (with EXP-NBD103 and SQK-LSK109) protocol (ONT).

### MinION library preparation

The barcoded pooled amplicon DNA was purified using Ampure XP magnetic beads. Each purified barcoded amplicon pool was quantified with Qubit, and pooled in equimolar quantities to generate the final library. The final 700ng DNA library was prepared for MinION sequencing as in the 1D Native barcoding DNA (with EXP-NBD103 and SQK-LSK109) protocol (ONT).

### MinION sequencing data analysis

The MinION sequencer was run for 1 hour 14 minutes, using an R9, 9.4 flowcell. Bases were called with guppy v2.3.7 (ONT) from the raw fast5 sequencing files from the MinION sequencer, followed by barcode demultiplexing also using guppy v2.3.7 (ONT), generating a fastq file for each barcode. Alternate basecalling was also performed using albacore v2.3.4 (ONT) and scrappie v1.4.0-0d933c7 (ONT), followed by barcode demultiplexing using guppy. Pool demultiplexing and alignment was performed for all fastq files using minimap2.

Alignment and coverage statistics were calculated using samtools 1.10 [18].

Variants were called and genotyped in the MinION sequencing data using nanopolish v0.13.2 and then compared with the Sanger generated data [19].

## Results

### Read characteristics

A single sequencing run of the MinION produced 1,096,297 reads corresponding to approximately 11,000 reads per sample. Of the total reads, 983,434 (89.7%) passed the quality qscore of seven, with a mean qscore of 9.7 and median qscore of 10.1. Guppy assigned the majority of reads (990,311) to the expected barcodes, with 93.0% of the correctly barcoded reads aligned to the reference by minimap2. The average sequence error rate was 3.05% for substitutions, 2.59% for deletions, and 1.73% for insertions generated by Alfred [20], consistent with expected error rates for the guppy basecaller [12, 21]. Metrics of the run are detailed in Table 1.

**Table 1.**
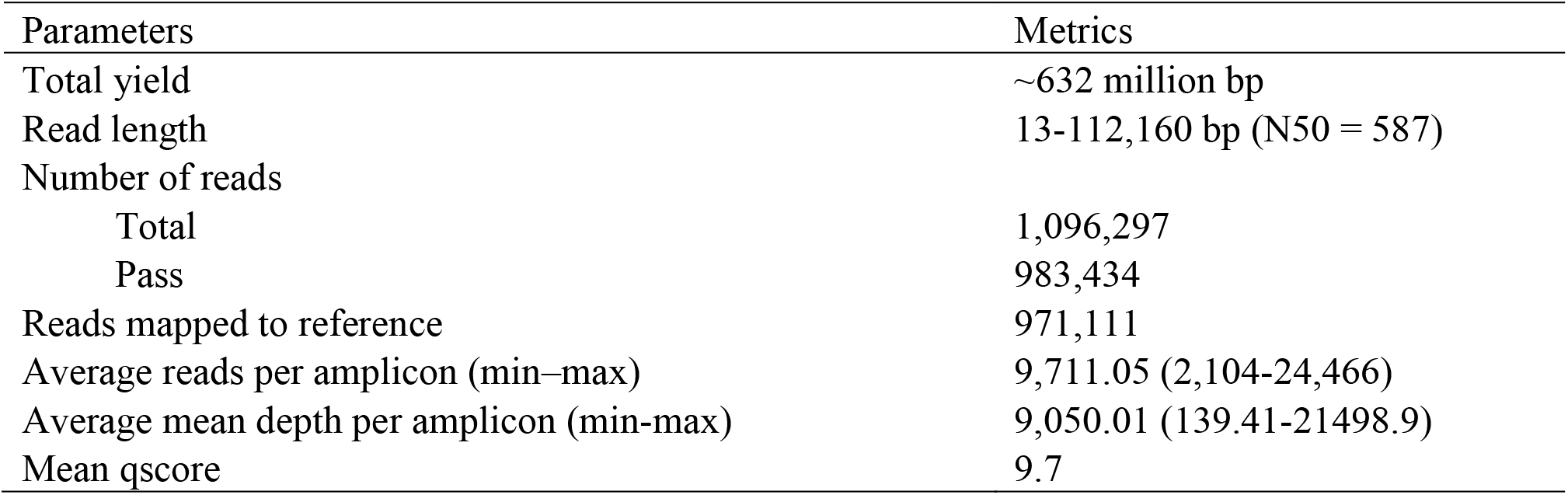
Metrics of the nanopore sequencing run

### Genotype validation

Sequences were basecalled by guppy and aligned to the appropriate reference by minimap2. Nanopolish called 29/30 of the variants of interest (97/100 PCR amplicons). The genotypes called by nanopolish were in concordance with the previously generated Sanger sequence. The variant which could not be identified by nanopolish was an insertion of an AT at the junction of two thymines and a homopolymer run of five cytosines (ref: CCCCCT, alt: CCCCC**CT**T; Supplementary Table 1).

The ability to call the correct number of bases in homopolymer runs is a known limitation of the Oxford Nanopore sequencing technology [8, 22]. Wick, Judd & Holt (2019) reported different levels of success for homopolymer calls from various basecallers [22]. Therefore, two alternative basecalling algorithms, albacore v2.3.4 (ONT) and scrappie v1.4.0-0d933c7 (ONT), were run in an attempt to identify the AT insertion. Two additional alignments were generated by minimap2 from the sequences generated from albacore and scrappie, however, neither could call the correct number of bases in the cytosine homopolymer run, or identify the homopolymer adjacent indel.

### Depth of coverage analysis

It was anticipated that pooling equimolar quantities for each PCR amplicon would result in comparable depth of coverage for each product when aligned to the respective reference. However, the average depth of sequence coverage for each amplicon widely varied between 21,498 and 139.41 fold (mean = 9,050, std err = 538.21). The majority of variance was between target regions rather than between PCR amplicons of the same target region from different individuals (Figure 1), with the average depth per target region varying significantly between 171x and 17740x [ANOVA: F(29, 70) = 10.37; P = 1.093×10^-15^].

**Figure 1.**
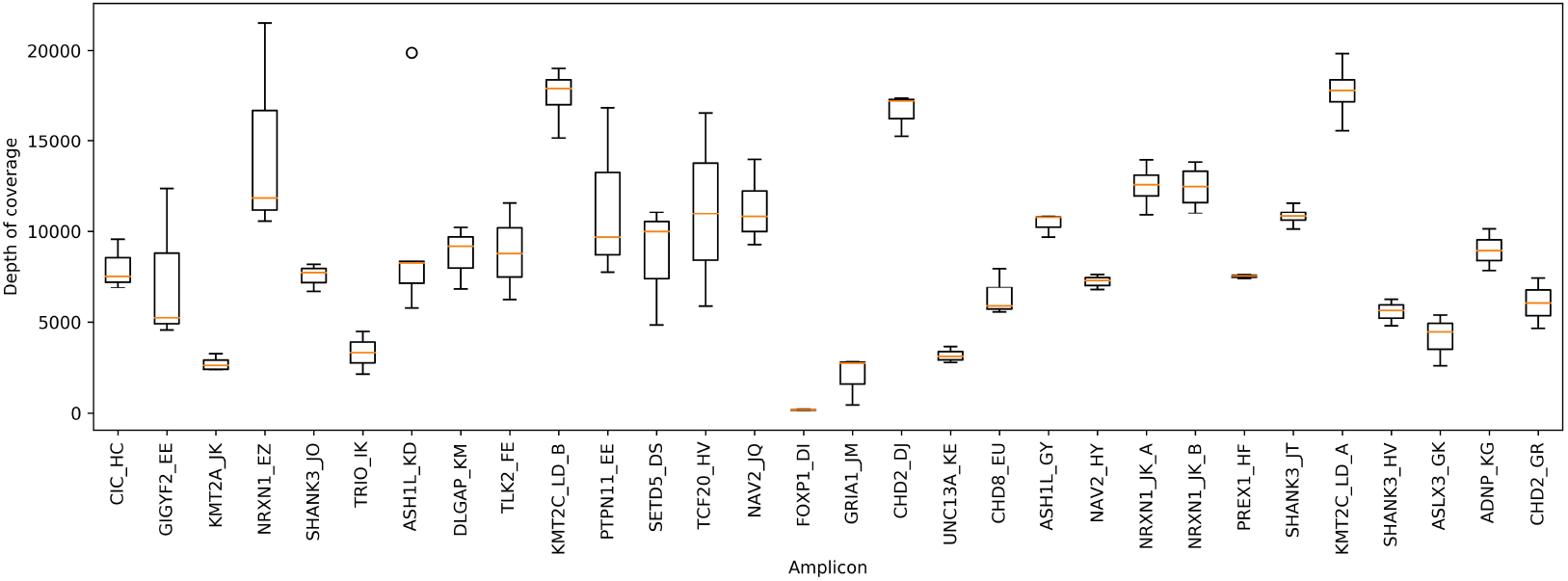
Depth of coverage of target regions. The name of each target region represents the gene name and the assigned pedigree ID. The average depth of sequence coverage varied significantly for different PCR amplicons, between 21,498 and 139.41 fold. The depth of coverage for PCR amplicons was significantly more uniform within target regions than between, with the average depth per target region varying significantly between 171x and 17740x [ANOVA: F(29, 70) = 10.37; P = 1.093×10-15]. A single outlier for the gene *ASH1L* is indicated by the circle.

There was a positive correlation between the nucleotide length of the PCR amplicon and the depth of coverage of the aligned sequence (Figure 2A). Conversely, there was a negative correlation between PCR amplicon GC content and depth of coverage (Figure 2B). While there was a negative correlation between PCR amplicon length and PCR amplicon GC content (Figure 2C), the coefficient of determination is low (R^2^ = 0.0526). This suggests that they independently affect the depth of sequence coverage. The lack of relationship was ratified by an insignificant interaction term in a linear regression (P=0.206).

**Figure 2.**
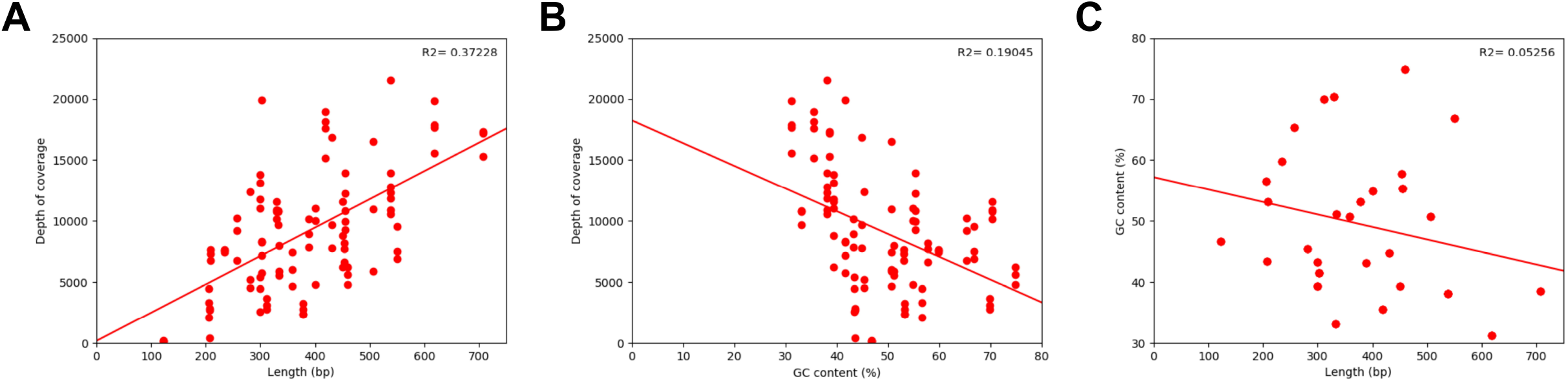
Alignment characteristics. **A.** Each PCR amplicon is plotted (100 total), demonstrating a positive relationship between the length of the product and depth of coverage. **B.** Each PCR amplicon is plotted (100 total), demonstrating a negative relationship between the GC content of the product and depth of coverage. **C.** Length of amplicon compared to the GC content of each different target region (30 total). The lack of significant relationship between these variables (R2=0.05256) indicates that each independently influences the depth of coverage of sequenced PCR amplicons

## Discussion

The rapid uptake of genome re-sequencing following the human genome project has dramatically improved our understanding of human disease. The discovery of pathogenic and disease susceptibility variants has contributed to a shift from phenotype-driven diagnosis to genotype-driven diagnosis. While there has been a substantial decrease in sequencing cost in recent years, the limited funding available for research and molecular diagnostic facilities often does not allow for WGS or WES of all individuals recruited or referred for genetic analysis. Thus, sequencing of targeted DNA intervals remains a common low cost alternative for applications such as mutational and inheritance confirmation [23], forensics [5] and targeted mutation screening for familial conditions [6].

In large studies of multiple individuals, targeted sequencing by traditional methods such as Sanger sequencing is still relatively costly at approximately $7 USD per reaction [24, 25]. We have described a method to genotype loci in up to 100 PCR amplicons across 82 individuals simultaneously using Oxford Nanopore Technology’s MinION sequencer. The method described in this report is comparable in cost to Sanger sequencing ($6.24 USD per PCR amplicon [26]), with opportunities to reduce the cost further (as discussed below). An additional benefit of MinION sequencing for targeted re-sequencing is the rapid return of results with the process from library preparation to analysed results achievable within a day.

Oxford Nanopore Technologies sequencing incorporates native barcoding, facilitating the sequencing of multiple samples simultaneously. By pooling PCR amplicons targeting multiple regions within the same barcode, we have expanded the utility of multiplexing with the MinION sequencing technology, reducing the per-amplicon cost of sequencing from $52 USD to $6.24 USD. The method we described is not specific to the target regions that we used for the analyses presented here, but can be easily modified to determine the presence of any variant by designing primers to any target of interest, provided the PCR amplicons to be pooled using a single barcode do not share significant similarity.

While this assay interrogated 100 PCR amplicons, the assay could be expanded, further increasing cost-effectiveness. Provided the reads can be correctly demultiplexed, the limit to the number of amplicons that can be interrogated is determined by sufficient reads for each amplicon and thus is a function of the yield of the flow cell and the length of the target regions of interest. Based on the error rate of 7.37% in this report, and successful genotyping from samples with 139x and 172x coverage, we estimate that coverage of 100x should be adequate for establishing an accurate genotype in the majority of cases. This estimate is consistent with a previous report of 80x depth sufficient to generate consensus sequence [27].

The reported maximum yield from a minION single flow cell is 50 Gbp [26], therefore with the metrics and amplicon size from this experiment, a theoretical 943,153 genetic loci could be accurately genotyped from a single flow cell with sufficient barcoding. However, this would require uniform depth of coverage for all amplicons and expending the full capacity of the flow cell over 72 hours. Liou, et al. also experienced an uneven distribution of reads with more than four-fold difference in depth between the lowest depth and the mean [17], and subsequently performed simulations to determine the maximum number of samples that could be genotyped in a single run. From a run of 6.3 million reads, consisting of 843 Mbp of sequence, with an average amplicon length of ~1300 bp, they determined a 95% accuracy in genotyping would be maintained with 7,000 amplicons. Given the estimated minimum depth required for genotyping was consistent between our results and those from Liou, et al., we expect that 7,000 amplicons could also be used as an upper limit for the method that we have described here. For example, this method (with adequate barcoding) would allow for the screening 70 loci in 96 individuals, while only using 1.7% of the maximum yield of the flow cell.

Expanding the number of amplicons targeting the same genetic region investigated in a single sequencing run can be supported by increasing the number of barcodes. This can either be through ONT native barcodes with a total of 96 available using the Native Barcoding Expansion 96 (EXP-NBD196) or through a dual barcode approach such as that used by Liou, et al., [17] and Srivathsan et al. [28]. The dual barcoding method applies barcoded PCR primers to incorporate barcodes into the products before pooling and addition of secondary ONT native barcodes. Furthermore, given the small size of the PCR amplicons typically used for targeted sequencing, the assay could be optimised for the cheaper ‘Flongle’ MinION flow cells capable of generating up to 2.8 Gbp of sequence. The Flongle would reduce the cost of sequencing to $2.39 USD per amplicon if sequencing 100 amplicons, or as little as 3 cents if sequencing 7,000 amplicons [29, 30].

Following decovolution, 29 of 30 of the investigated genetic loci were successfully genotyped. Nanopolish identified 28 single nucleotide subtitutions and one single base pair insertion from basecalls by guppy. The variant that was not identified during variant calling by any of the algorthims tested here is an insertion of adenosine and thymine at the junction of two thymines and a homopolymer run of five cytosines. The difficulty of basecalling MinION sequence surrounding stretches of single bases is a well-reported limitation of the technology [9, 28, 31].

Nanopore sequencing utilises nanoscale pores for which changes in the ionic current passing through the pore changes as DNA/RNA passes through. This change in current is used to determine the bases present in the pore and thus nucleotide sequence of the template [8]. Consequently, a repeatedly reported limitation of the technology is the difficulty to correctly identify the length of homopolymer runs greater than the k-mer of the nucleotides determining the signal from the pore as these extended homopolymer runs do not result in a change in the current through the pore [8]. For the flowcells used in this experiment (R9, 9.4) the signal is primarily determined from the three central nucleotides within the pore [32], thus accuractely determining the length of a homopolymer greater than as few as four nucleotides is technically challenging. Similar to the increase in raw read accuracy that has been achieved with updates in chemistry and software tools [8], we anticipate improvements to homopolymer calls. Particularly, ONT reports better homopolymer identification with R10, 10.3 pores released in January 2020 [33].

Oikonomopoulos, et al. (2016) reported that the proportion of MinION reads very closely mirrors the number of template molecules in the library [31]. Therefore, we presumed including equimolar quantities of PCR amplicons would result in an equal depth of coverage for each product. However, we observed significant variance between target regions, with less variance between PCR amplicons from the same target. Therefore, inherent differences between the amplicons resulted in differences in read depth. Specifically, within the bounds of amplicons investigated here, MinION sequencing has a bias for longer amplicon length and lower GC content.

There are mixed reports of GC content and length bias in MinION sequencing. Oikonomopoulos, et al. (2016) did not observe any length or GC content bias from the pooled library [31]. However, Li, et al. (2020) performed a direct comparison of the reads generated by MinION and Pacbio Sequel and Ion Torrent PGM and observed a significantly reduced number of reads with > 45% GC content from MinION relative to the other technologies [34]. Notably, there was a negative relationship between read counts and GC content > 40%. Both Li, et al. (2020) and Karlsson, et al. (2015) reported a normal distribution of the proportion of reads centred around ~7 kb [34, 35]. This distribution replicates the positive relationship that was observed in this report, if limited to the 122-707 bp window that was investigated here.

The lack of relationship between target sequence length and GC content indicated that the two variables independently affect depth of coverage. Therefore, pooling equimolar quantities of PCR amplicons may not be the best approach to achieve equal depth of coverage for all templates required for accurate variant identification while maximising the use of the flow cell. With a significantly larger number of sequencing runs targeting additional varied template sequences, a regression model incorporating depth of coverage, length, GC content and molarity may reveal the relationship between these variables and determine the optimum quantity of products to be pooled. Such a model will optimise the depth of coverage for each PCR amplicon, thus allowing for screening of the maximum number of amplicons in a single sequencing run.

## Conclusion

Oxford Nanopore Technology’s MinION sequencer allows for rapid and cost-effective development of variant discovery assays. Genotypes were called with a 97% accuracy, the only exception being an insertion adjacent to a homopolymer run. The difficulty to correctly call the number of bases in a homopolymer run is a known limitation of the technology which will likely be mitigated through improvements in chemistry and flow cells by ONT and the development of superior basecalling software. The depth of coverage does not scale with the number of molecules of DNA pooled, however the depth was sufficient for successful variant calling in 29/30 of the genetic variants of interest. Therefore, ONT MinION can be used as a time and cost-effective alternative to Sanger sequencing for large-scale variant identification from any short DNA template with adequate barcoding and read depth.

## Declarations

### Ethics approval and consent to participate

The genetic analysis and de-identified publication of variants for use cases was performed under the approval of the New Zealand Northern B Health and Disability Ethics Committee (12/NTB/59) in accordance with guidelines and regulations in the Ethical Guidelines for Observational Studies from the New Zealand National Ethics Advisory Committee. Parents and affected offspring provided written informed consent (with parents providing consent on behalf of their children, where appropriate).

### Competing interests

The authors declare that they have no competing interests.

### Funding

WW is supported by IHC Foundation, and The Kate Edger Educational Charitable Trust. The research was funded by Cure Kids, the Minds for Minds Charitable Trust, and the IHC Foundation.

### Authors’ contributions

WW performed MinION sequencing, completed downstream analysis, and composed the manuscript. WW and MJG conceptualised the statistical analyses. KL performed WES alignment and annotation. WW, VH, KM and JJ performed variant interpretation and generated PCR amplicons for sequencing. WW, JCJ and RGS conceptualised the experimental methods. JCJ and RGS critically reviewed the manuscript. All authors read and approved the final manuscript.

## Acknowledgements

The authors wish to thank Kristine Boxen the Genomics Centre, Auckland Science Analytical Services, The University of Auckland, Auckland, New Zealand for assistance with Sanger sequencing.

The authors also wish to acknowledge the contribution of NeSI high-performance computing facilities to the results of this research. NZ’s national facilities are provided by the NZ eScience Infrastructure and funded jointly by NeSI’s collaborator institutions and through the Ministry of Business, Innovation & Employment’s Research Infrastructure programme. URL https://www.nesi.org.nz.

## Notes

### Competing Interest Statement

The authors have declared no competing interest.

## References

1. Whitford W, Hawkins I, Glamuzina E, Wilson F, Marshall A, Ashton F, et al. Compound heterozygous SLC19A3 mutations further refine the critical promoter region for biotin-thiamine-responsive basal ganglia disease. 2017;3. doi:10.1101/mcs.a001909.

2. Jacobsen JC, Whitford W, Swan B, Taylor J, Love DR, Hill R, et al. Compound Heterozygous Inheritance of Mutations in Coenzyme Q8A Results in Autosomal Recessive Cerebellar Ataxia and Coenzyme Q10 Deficiency in a Female Sib-Pair. Springer, Berlin, Heidelberg; 2017. p. 31–6. doi:10.1007/8904_2017_73.

3. McRae JF, Clayton S, Fitzgerald TW, Kaplanis J, Prigmore E, Rajan D, et al. Prevalence and architecture of de novo mutations in developmental disorders. Nature. 2017;542:433–8.

4. Pfundt R, del Rosario M, Vissers LELM, Kwint MP, Janssen IM, de Leeuw N, et al. Detection of clinically relevant copy-number variants by exome sequencing in a large cohort of genetic disorders. Genet Med. 2017;19:667–75. doi:10.1038/gim.2016.163.

5. Sullivan W, Evans DG, Newman WG, Ramsden SC, Scheffer H, Payne K. Developing National Guidance on Genetic Testing for Breast Cancer Predisposition: The Role of Economic Evidence? Genet Test Mol Biomarkers. 2012;16:580–91. doi:10.1089/gtmb.2011.0236.

6. Jain M, Olsen HE, Paten B, Akeson M. The Oxford Nanopore MinION: delivery of nanopore sequencing to the genomics community. Genome Biol. 2016;17:239. doi:10.1186/s13059-016-1103-0.

7. Tyler AD, Mataseje L, Urfano CJ, Schmidt L, Antonation KS, Mulvey MR, et al. Evaluation of Oxford Nanopore’s MinION Sequencing Device for Microbial Whole Genome Sequencing Applications. Sci Rep. 2018;8:1–12.

8. Rang FJ, Kloosterman WP, de Ridder J. From squiggle to basepair: Computational approaches for improving nanopore sequencing read accuracy. Genome Biology. 2018;19:90. doi:10.1186/s13059-018-1462-9.

9. Cornelis S, Gansemans Y, Deleye L, Deforce D, Van Nieuwerburgh F. Forensic SNP Genotyping using Nanopore MinION Sequencing. Sci Rep. 2017;7:1–5.

10. Cornelis S, Gansemans Y, Vander Plaetsen AS, Weymaere J, Willems S, Deforce D, et al. Forensic tri-allelic SNP genotyping using nanopore sequencing. Forensic Sci Int Genet. 2019;38:204–10.

11. Leija-Salazar M, Sedlazeck FJ, Toffoli M, Mullin S, Mokretar K, Athanasopoulou M, et al. Evaluation of the detection of GBA missense mutations and other variants using the Oxford Nanopore MinION. Mol Genet Genomic Med. 2019;7. doi:10.1002/mgg3.564.

12. Liau Y, Cree SL, Maggo S, Miller AL, Pearson JF, Gladding PA, et al. A multiplex pharmacogenetics assay using the MinION nanopore sequencing device. Pharmacogenet Genomics. 2019;29:207–15.

13. Quick J, Grubaugh ND, Pullan ST, Claro IM, Smith AD, Gangavarapu K, et al. Multiplex PCR method for MinION and Illumina sequencing of Zika and other virus genomes directly from clinical samples. Nat Protoc. 2017;12:1261–6. doi:10.1038/nprot.2017.066.

14. Freed NE, Vlková M, Faisal MB, Silander OK. Rapid and Inexpensive Whole-Genome Sequencing of SARS-CoV2 using 1200 bp Tiled Amplicons and Oxford Nanopore Rapid Barcoding. doi:10.1101/2020.05.28.122648.

15. Geoghegan JL, Ren X, Storey M, Hadfield J, Jelley L, Jefferies S, et al. Genomic epidemiology reveals transmission patterns and dynamics of SARS-CoV-2 in Aotearoa New Zealand. Nat Commun. 2020;11:1–7. doi:10.1038/s41467-020-20235-8.

16. Li J, Wang H, Mao L, Yu H, Yu X, Sun Z, et al. Rapid genomic characterization of SARS-CoV-2 viruses from clinical specimens using nanopore sequencing. Sci Reports 2020 101. 2020;10:1–10. doi:10.1038/s41598-020-74656-y.

17. Liou CH, Wu HC, Liao YC, Lauderdale TLY, Huang IW, Chen FJ. Nanomlst: Accurate multilocus sequence typing using oxford nanopore technologies minion with a dual-barcode approach to multiplex large numbers of samples. Microb Genomics. 2020;6:1–8. doi:10.1099/mgen.0.000336.

18. Li H, Handsaker B, Wysoker A, Fennell T, Ruan J, Homer N, et al. The Sequence Alignment/Map format and SAMtools. Bioinformatics. 2009;25:2078–9. doi:10.1093/bioinformatics/btp352.

19. Loman NJ, Quick J, Simpson JT. A complete bacterial genome assembled de novo using only nanopore sequencing data. Nat Methods. 2015;12:733–5.

20. Rausch T, Hsi-Yang Fritz M, Korbel JO, Benes V. Alfred: Interactive multi-sample BAM alignment statistics, feature counting and feature annotation for long- and short-read sequencing. Bioinformatics. 2019.

21. De Coster W, De Rijk P, De Roeck A, De Pooter T, D’Hert S, Strazisar M, et al. Structural variants identified by Oxford Nanopore PromethION sequencing of the human genome. Genome Res. 2019;29:1178–87. doi:10.1101/gr.244939.118.

22. Wick RR, Judd LM, Holt KE. Performance of neural network basecalling tools for Oxford Nanopore sequencing. Genome Biol. 2019;20:129. doi:10.1186/s13059-019-1727-y.

23. Jacobsen JC, Whitford W, Swan B, Taylor J, Love DR, Hill R, et al. Compound heterozygous inheritance of mutations in Coenzyme Q8A results in autosomal recessive cerebellar ataxia and coenzyme Q10 deficiency in a female sib-pair. In: JIMD Reports. Springer; 2018. p. 31–6.

24. Sanger Sequencing Service Fees | Center for Genome Research and Biocomputing | Oregon State University. https://cgrb.oregonstate.edu/core/sanger-sequencing/sanger-sequencing-service-fees. Accessed 21 Jun 2021.

25. Sanger Pricing | Department of Biochemistry. https://www.bioc.cam.ac.uk/dnasequencing/sanger-sequencing/pricing. Accessed 21 Jun 2021.

26. MinION | Oxford Nanopore Technologies. https://nanoporetech.com/products/minion. Accessed 17 May 2021.

27. Liao YC, Cheng HW, Wu HC, Kuo SC, Lauderdale TLY, Chen FJ. Completing Circular Bacterial Genomes With Assembly Complexity by Using a Sampling Strategy From a Single MinION Run With Barcoding. Front Microbiol. 2019;10:2068. doi:10.3389/fmicb.2019.02068.

28. Srivathsan A, Baloğlu B, Wang W, Tan WX, Bertrand D, Ng AHQ, et al. A MinION™-based pipeline for fast and cost-effective DNA barcoding. Mol Ecol Resour. 2018;18:1035–49. doi:10.1111/1755-0998.12890.

29. Nanopore Store https://store.nanoporetech.com/native-barcoding-expansion-1-12.html. Accessed 1 Jun 2021.

30. Product comparison | Oxford Nanopore Technologies. https://nanoporetech.com/products/comparison. Accessed 1 Jun 2021.

31. Oikonomopoulos S, Wang YC, Djambazian H, Badescu D, Ragoussis J. Benchmarking of the Oxford Nanopore MinION sequencing for quantitative and qualitative assessment of cDNA populations. Sci Rep. 2016;6. doi:10.1038/srep31602.

32. Brown C. Oxford Nanopore Technologies: Owl stretching with examples. 2016. https://www.youtube.com/watch?v=JmncdnQgaIE. Accessed 4 Aug 2020.

33. R10.3: the newest nanopore for high accuracy nanopore sequencing - now available in store. https://nanoporetech.com/about-us/news/r103-newest-nanopore-high-accuracy-nanopore-sequencing-now-available-store. Accessed 17 May 2021.

34. Li Y, He X zhou, Li M hui, Li B, Yang M jie, Xie Y, et al. Comparison of third-generation sequencing approaches to identify viral pathogens under public health emergency conditions. Virus Genes. 2020;56:288–97. doi:10.1007/s11262-020-01746-4.

35. Karlsson E, Lärkeryd A, Sjödin A, Forsman M, Stenberg P. Scaffolding of a bacterial genome using MinION nanopore sequencing. Sci Rep. 2015;5:1–8. doi:10.1038/srep11996.

