## Supplemental Methods for "Optimised multiplex amplicon sequencing for mutation identification using the MinION nanopore sequencer"

**Supplementary methods**

**DNA extraction**

Whole exome sequencing was performed using the SureSelect XT Human All Exon v6 (Agilent, Santa Clara, CA, USA) and Illumina Novaseq 6000 by Macrogen to obtain 59 million 100bp paired-end reads for each individual. Reads were mapped to the human GRCh37.p13 reference using bwa (v 0.7.4) [1], duplicates marked with Picard (v 1.96) [2] followed by indel realignment and base quality score recalibration using the Genome Analysis Toolkit (GATK, v 3.4-46). SNP and indel variants were identified using GATK HaplotypeCaller, and GenotypeGVCFs as described in Van der Auwera et al [3].

Genetic variants for this report were selected from a study focussing on the identification of causative genetic variants for individuals diagnosed with autism spectrum disorder (ASD). Variants were therefore prioritised if they resided in genes expressed in the brain [4], had previously been reported as causative in ASD or neurodevelopmental disorders [5], and by SFARI score (a scoring system based on the strength of evidence supporting the role of the gene to ASD risk [6]). Additionally, variants were filtered to prioritise those observed at a minor allele frequency of less than 0.1% in the 1000 Genomes [7], HapMap [8], genome aggregation database (gnomAD, [9]), or our in-house dataset.

**Supplementary Table 1.** Amplicons Sequenced

| **Amplicon** | **Primers** | **Product size (bp)** | **GC content (%)** | **Variant** | **Individuals sequenced by minION** | **WES genotype; allele depth** | **MinION genotype; strand support** |
| --- | --- | --- | --- | --- | --- | --- | --- |
| CIC_HC | Primer F  5’ GTTGAAGATCCGTGAGGTGC 3’  Primer R  5’ TCACACGCTCCAGGTTATGT 3’ | 551 | 66.79 | ENST00000575354.2:c.4718C>T | Proband^†^  Mother  Father | 0/1; 11,12 | 0/1; 2014,1435,2199,1736  0/1; 2438,1804,3505,2419  0/0‡ |
| GIGYF2_EE | Primer F  5’ TCAGTCCATTTGAGTTTGCGG 3’  Primer R  5’ TCCTCTAAGCACCATTCGGG 3’ | 282 | 45.39 | ENST00000409451.3:c.817C>T | Proband^†^  Mother  Father | 0/1; 28,39 | 0/1; 1205,1131,1148,1266  0/0‡  0/1; 1335,1393,1289,1408 |
| KMT2A_JK | Primer F  5’ ACGTGGTGGACTCTAGTCAGA 3’  Primer R  5’ GCTGTTTGAGACATCAGTGCT 3’ | 378 | 53.17 | ENST00000534358.1:c.3974G>A | Proband^†^  Sibling^†^  Mother  Father | 0/1; 28,35  0/0^‡^ | 0/1; 397,1118,448,1012  0/0‡  0/0‡  0/1; 379,941,375,826 |
| NRXN1_EZ | Primer F  5’ CATGTGAAGGGAGACCGTGT 3’  Primer R  5’ AGCGCGTGGTGAAAGATATTG 3’ | 538 | 38.10 | ENST00000404971.1:c.3071A>G | Proband^†^  Mother  Father | 0/1; 33,25 | 0/1; 2645,2884,2610,2755  0/1; 5433,6040,5233,5387  0/0‡ |
| SHANK3_JO | Primer F  5’ CAGGTGAGACCTGAGCGTG 3’  Primer R  5’ CAACACCAAATACCCCTCGC 3’ | 454 | 57.71 | ENST00000262795.3:c.898C>T | Proband^†^  Mother  Father | 0/1; 60,50 | 0/1; 1456,1971,1642,1906  0/0‡  0/1; 2002,2291,2045,2176 |
| TRIO_IK | Primer F  5’ GGTCCTATCAATCTGTCGGGG 3’  Primer R  5’ TCAGGGCCCTTCCAGGTAAT 3’ | 205 | 56.59 | ENST00000344204.4:c.8066T>C | Proband^†^  Mother  Father | 0/1; 62,48 | 0/1; 1068,612,1113,601  0/0‡  0/1; 733,298,816,304 |
| ASH1L_KD | Primer F  5’ TGACCTATGACCAACGTTCAAGT 3’  Primer R  5’ TCAAAGCATGAAAAGCAGCCTC 3’ | 303 | 41.58 | ENST00000392403.3:c.1731G>C | Proband^†^  Sibling1^†^  Sibling2^†^  Mother  Father | 0/1; 31,29  0/1; 37,28  0/1; 47,25 | 0/1; 2195,1600,2005,1674  0/1; 2747,1719,2486,1739  0/1; 2043,1103,1694,1150  0/1; 6119,4287,5795,4346  0/0‡ |
| DLGAP_KM | Primer F  5’ ACACTCGTCCTTCAGCTCTTG 3’  Primer R  5’ CCATGAAAGGGCTATCAGGCA 3’ | 257 | 65.37 | ENST00000315677.3:c.85C>T | Proband^†^  Mother  Father | 0/1; 20,15 | 0/1; 1565,1819,1961,1599  0/0‡  0/1; 2223,2662,3036,2425 |
| TLK2_FE | Primer F  5’ GAGGACTCTCCCTGAGTATCCA 3’  Primer R  5’ TAATGATTTGGGGAGAAAGTCTGC 3’ | 450 | 39.33 | ENST00000346027.5:c.1784C>G | Proband^†^  Mother  Father | 0/1; 22,21 | 0/1; 1980,2621,1948,2534  0/0‡  0/0‡ |
| KMT2C_LD_B | Primer F  5’ AATGGTGAGTCAGAGTATCCCA 3’  Primer R  5’ AGTTATCGCTTAAAGCAGTTGAATA 3’ | 419 | 35.56 | ENST00000262189.6:c.467C>T | Proband^†^  Sibling1^†^  Mother  Father | 0/1; 28,32  0/1; 30,37 | 0/1; 4595,4882,4049,4458  0/1; 4867,4936,4689,4905  0/1; 4634,4978,4301,4588  0/0‡ |
| PTPN11_EE | Primer F  5’ TCCTGACTTCTGCCACTTCGT 3’  Primer R  5’ CAAAAGGAGAGCGTATCCAAGAGG 3’ | 431 | 44.78 | ENST00000351677.2:c.1492C>T | Proband^†^  Mother  Father | 0/1; 20,24 | 0/1; 3872,4798,3536,5104  0/0‡  0/1; 1517,2595,1285,2677 |
| SETD5_DS | Primer F  5’ GGGACTTGTTCGCGTCCTTAT 3’  Primer R  5’ TCTGAGGTTGGCGAGTCTGA 3’ | 401 | 54.86 | ENST00000402198.1:c.3929C>T | Proband^†^  Mother  Father | 0/1; 43,58 | 0/1; 1480,1056,1478,1030  0/1; 3480,2370,3446,2281  0/0‡ |
| TCF20_HV | Primer F  5’ CAGTCGCTTTTCTGGTACCCC 3’  Primer R  5’ AATGCACAGGCTTATGGAACAC 3’ | 507 | 50.69 | ENST00000359486.3:c.1105A>G | Proband^†^  Mother  Father | 0/1; 79,71 | 0/1; 1507,1566,1477,1531  0/0‡  0/1; 4628,4444,4033,4058 |
| NAV2_JQ | Primer F  5’ TAGCACCTCGAACTGTCTGC 3’  Primer R  5’ GAGGCTCTGCATCATACCCA 3’ | 455 | 55.38 | ENST00000396087.3:c.7261G>A | Proband^†^  Sibling1^†^  Sibling2^†^  Mother  Father | 0/1; 65,47  0/1; 72,61  0/1; 55,53 | 0/1; 2775,2551,1985,2342  0/1; 2943,2740,2105,2633  0/1; 3871,3760,3168,3700  0/0‡  0/1; 3080,2913,2269,2968 |
| FOXP1_DI | Primer F  5’ CTGAGAAAGCTTACCTTCCACG 3’  Primer R  5’ TCACAGGCCATTCTCGAATCT 3’ | 122 | 46.7 | ENST00000491238.1:c.1472A>G | Proband^†^  Mother  Father | 0/1; 40,36 | 0/1; 41,28,34,32  0/1; 33,23,27,17  0/0‡ |
| GRIA1_JM | Primer F  5’ CTTTGGTCCGGGAAGAAGTT 3’  Primer R  5’ ATTCATAGGGACTGAAGCGGC 3’ | 207 | 43.49 | ENST00000518783.1:c.1568G>A | Proband^†^  Mother  Father | 0/1; 28,12 | 0/1; 118,100,96,106  0/0‡  0/0‡ |
| **CHD2_DJ** | **Primer F**  **5’ AGGCTCTTGCCAAAGGAACA 3’**  **Primer R**  **5’ CCAGTGTAGGAAGGTTGGGG 3’** | **707** | **38.61** | **ENST00000394196.4:c.2423_2424insAT** | **Proband^†^**  **Mother**  **Father** | **0/1; 49,31** | **0/0‡**  **0/0‡**  **0/0‡** |
| UNC13A_KE | Primer F  5’ ATCTTGGTTCAGCACCGGG 3’  Primer R  5’ GAGTATTGCAGGGAGGCGTT 3’ | 312 | 69.87 | ENST00000428389.2:c.205G>A | Proband^†^  Mother  Father | 0/1; 16,18 | 0/1; 1487,333,1681,338  0/1; 1250,364,1315,321  0/0‡ |
| CHD8_EU | Primer F  5’ CACAATGCCAGCCGATCTTC 3’  Primer R  5’ TCCGCACTTTTGCTCGACT 3’ | 334 | 51.20 | ENST00000399982.2:c.5665C>T | Proband^†^  Mother  Father | 0/1; 34,36 | 0/1; 2685,1142,1204,1067  0/1; 2292,1045,1445,986  0/0‡ |
| ASH1L_GY | Primer F  5’ TATCACTGGTGTGCTTTACTTCCT 3’  Primer R  5’ GTTTCACTTGCCAGCATTTTTGC 3’ | 332 | 33.13 | ENST00000392403.3:c.7061T>G | Proband^†^  Mother  Father | 0/1; 25,25 | 0/1; 2319,3145,2211,2407  0/1; 2653,3480,2358,2653  0/0‡ |
| NAV2_HY | Primer F  5’ GACAAGTGTGTCTGCTTCGG 3’  Primer R  5’ AGCAGGTATTGCGTGGAAGG 3’ | 209 | 53.11 | ENST00000396087.3:c.2153G>A | Proband^†^  Mother  Father | 0/1; 84,73 | 0/1; 1456,2057,1438,1889  0/1; 1504,2440,1518,2340  0/0‡ |
| NRXN1_JK_A | Primer F  5’ CATGTGAAGGGAGACCGTGT 3’  Primer R  5’ AGCGCGTGGTGAAAGATATTG 3’ | 538 | 38.10 | ENST00000404971.1:c.3113G>A | Proband^†^  Sibling1^†^  Mother  Father | 0/0‡  0/1; 91,8^*^ | 0/0‡  0/0‡  0/0‡  0/0‡ |
| NRXN1_JK_B | Primer F  5’ ATGTGTTGATTGCCTTGCTTTGA 3’  Primer R  5’ GGTAGTGGAGGCCAAGTTGTAA 3’ | 300 | 39.33 | ENST00000404971.1:c.780A>T | Proband^†^  Sibling1^†^  Mother  Father | 0/1; 24,22  0/0‡ | 0/1; 2961,2688,3006,2733  0/0‡  0/1; 3148,2943,3118,2953  0/0‡ |
| PREX1_HF | Primer F  5’ CCTTGGACTGGCTATCCCCT 3’  Primer R  5’ ACTTTGGGTTCCTCCTGTCA 3’ | 234 | 59.83 | ENST00000371941.3:c.3727C>T | Proband^†^  Mother  Father | 0/1; 104,66 | 0/1; 1991,1896,1976,1611  0/1; 2034,2071,1875,1657  0/0‡ |
| SHANK3_JT | Primer F  5’ AGTCACCCGAGGACAAGAAGT 3’  Primer R  5’ CCTCATCGCTGGACGACAG 3’ | 330 | 70.30 | ENST00000262795.3:c.3700G>A | Proband^†^  Sibling1^†^  Mother  Father | 0/1; 53,33  0/1; 42,25 | 0/1; 3729,2557,1539,2648  0/1; 4183,2898,1778,3074  0/0‡  0/1; 3941,2874,1670,2714 |
| KMT2C_LD_A | Primer F  5’ CCCACTTTATTGAAAGACTACAGGT 3’  Primer R  5’ AATCTTTCTTGTGAGGTCTAGTTGT 3’ | 619 | 31.18 | ENST00000262189.6:c.1759_1769del | Proband^†^  Sibling1^†^  Mother  Father | 0/0‡  0/1; 41,24 | 0/0‡  0/1; 5285,5060,5305,4889  0/0‡  0/0‡ |
| SHANK3_HV | Primer F  5’ CCGGGTGGCCTCGACT 3’  Primer R  5’ GTACATCCACAAACAGGGGTC 3’ | 460 | 74.78 | ENST00000262795.3:c.3227C>G | Proband^†^  Mother  Father | 0/1; 40,27 | 0/1; 1326,1929,1392,1891  0/1; 916,1528,1056,1564  0/0‡ |
| ASLX3_GK | Primer F  5’ TCCTAATGTCTGTTGACAGTGCAAA 3’  Primer R  5’ TTGGGGGAATCAAGACAGAGCTA 3’ | 300 | 43.33 | ENST00000269197.5:c.3889C>T | Proband^†^  Mother  Father | 0/1; 29,32 | 0/1; 1947,1085,1725,897  0/0‡  0/1; 970,439,884,414 |
| ADNP_KG | Primer F  5’ GCGGCCATCTTTTCCACATC 3’  Primer R  5’ TCTTCTCTCTCATCGGGCCA 3’ | 389 | 43.19 | ENST00000396029.3:c.1490A>G | Proband^†^  Mother  Father | 0/1; 32,41 | 0/1; 1342,3068,1230,3670  0/0‡  0/0‡ |
| CHD2_GR | Primer F  5’ TGCAGTCATCAGATCATTCTTTCT 3’  Primer R  5’ TCATGTATACGTCACTTTCTCCCT 3’ | 359 | 50.70 | ENST00000394196.4:c.5032C>T | Proband^†^  Mother  Father | 0/1; 91,68 | 0/1; 1937,1725,1939,2118  0/0‡  0/1; 1124,1203,1120,1396 |

Amplicon names represent the gene for which the locus of interest is located followed by the assigned pedigree ID

† Individuals for which whole exome sequence was available

*Variant determined to be homozygous reference by Sanger sequencing

‡Variants not called by variant caller are indicated by 0/0

Standard annotations of 0 for the reference allele, 1 for the first alternate, 0/0 for homozygous refence, and 0/1 for heterozygous variants.

The bolded row represents the variant that was unable to be correctly genotyped from aligned MinION sequence

**References**

1. Li H, Durbin R. Fast and accurate short read alignment with Burrows-Wheeler transform. Bioinformatics. 2009;25:1754–60. doi:10.1093/bioinformatics/btp324.

2. BroadInstitute. Picard Tools - By Broad Institute. 2016. http://broadinstitute.github.io/picard/. Accessed 3 Sep 2018.

3. Van der Auwera GA, Carneiro MO, Hartl C, Poplin R, del Angel G, Levy-Moonshine A, et al. From FastQ Data to High-Confidence Variant Calls: The Genome Analysis Toolkit Best Practices Pipeline. In: Current Protocols in Bioinformatics. Hoboken, NJ, USA: John Wiley & Sons, Inc.; 2013. p. 11.10.1-11.10.33. doi:10.1002/0471250953.bi1110s43.

4. GTEx Consortium. The Genotype-Tissue Expression (GTEx) pilot analysis: multitissue gene regulation in humans. Science (80- ). 2015;348:648–60. doi:10.1126/science.1262110.

5. McKusick-Nathans Institute of Genetic Medicine, Johns Hopkins University (Baltimore M. Online Mendelian Inheritance in Man, OMIM®. https://www.omim.org/. Accessed 4 Mar 2020.

6. Abrahams BS, Arking DE, Campbell DB, Mefford HC, Morrow EM, Weiss LA, et al. SFARI Gene 2.0: a community-driven knowledgebase for the autism spectrum disorders (ASDs). Mol Autism. 2013;4:36. doi:10.1186/2040-2392-4-36.

7. The 1000 Genomes Project Consortium. A global reference for human genetic variation. Nature. 2015;526:68–74. doi:10.1038/nature15393.

8. The International Hapmap Consortium. The International HapMap Project. Nature. 2003;426:789–96. doi:10.1038/nature02168.

9. Lek M, Karczewski KJ, Samocha KE, Banks E, Fennell T, O’Donnell-Luria AH, et al. Analysis of protein-coding genetic variation in 60,706 humans. Nature. 2016;536:285–91. doi:10.1101/030338.
